# Evidence for selective attention in the insect brain

**DOI:** 10.1101/041889

**Authors:** Benjamin L. de Bivort, Bruno van Swinderen

## Abstract

The capacity for selective attention appears to be required for any animal responding to an environment containing multiple objects, although this has been difficult to study in smaller animals such as insects. Clear operational characteristics of attention however make study of this crucial brain function accessible to any animal model. Whereas earlier approaches have relied on freely behaving paradigms placed in an ecologically relevant context, recent tethered preparations have focused on brain imaging and electrophysiology in virtual reality environments. Insight into brain activity during attention-like behavior has revealed key elements of attention in the insect brain. Surprisingly, a variety of brain structures appear to be involved, suggesting that even in the smallest brains attention might involve widespread coordination of neural activity.

**Highlights:**

- Insect behavior shows attention-like filtering and allows concurrent neural recording
- Recent work finds compelling correlates of visual attention in the central complex
- Putative goal-directed stimulus-filtering is present in neuropils across the brain

## Introduction

Do insects have selective attention? Do they share our ability to focus on only essential information from the environment at any given time, while actively ignoring a multitude of other potentially distracting stimuli [1,2]? Although this may seem difficult to address in any animal incapable of self-report, visual attention has been studied for a long time in animals ranging from primates [3,4] to honeybees [5,6]. Recently, new techniques for simultaneously tracking behavior and brain activity in insects are providing some pertinent insights. Few insect studies, however, have directly tested for visual attention-like processes, and these insights, mostly measuring fly and bee vision, are primarily placed in a sensory processing context. To investigate insect attention more directly, much can of course be learned from other animal studies [7]. Although a great variety of attention experiments have been designed for different sensory modalities, these are often structured in a similar way. Most experiments probe one of four operational characteristics of attentional processes (Box 1), and all of these have been applied to insects in some way: 1. Attention promotes responsiveness to a subset of incoming information at any given time, whether these are a common feature (e.g., a color [6]), or individual objects (e.g., a moving bar [8]), 2. Attention is a limited resource, meaning that more difficult tasks will make it less likely to notice other events [9], 3. Attention alternates serially among percepts with a characteristic timescale, meaning that individuals have a typical ‘attention span’ in a given environment [10]. This temporal aspect of attention has been linked to working memory in some studies [11]. 4. Crucially, one should be able to identify neural correlates for all of the above. Here, we focus on reviewing evidence for neural correlates of attention in the insect brain, and present a framework for further inquiry that requires moving beyond the usual ways in which insect brains are typically studied.

#### Box 1 - Operational characteristics of attentional phenomena

- <u>Unity</u> – Attention can be directed to a subset of stimulus features at any given time. *Test with distractor stimuli*.
- <u>Resource limitation</u> – Total attentional filtering capacity is limited. *Test with challenging tasks, i.e., high attentional loads*.
- <u>Alternation</u> – Attention shifts between percepts on characteristic timescales. *Test with cueing, training, or top-down instructions that probe fixation timescales*.
- <u>Neural correlates</u> – Attentional phenomena reflect patterns of neural activity. *Test with simultaneous physiological and behavioral recording*.

Each of the attention criteria discussed (Box 1) can be operationalized and framed in testable hypotheses. Brain activity readouts are especially valuable, because attention-like processes should in principle persist in the absence of correlated behavior or stimuli [12]. A typical human visual attention experiment, for example, will require subjects to fixate on a task of varying difficulty while target objects flash on or off around the fixation point. Task difficulty probes ‘attention load’, i.e., resource limitation. At the same time, attention can be redirected by competing objects. These distractors reduce performance by drawing attention away from the fixated task, thus probing the unity of attention and the timescales of attentional alternation. Readouts for human attention studies are diverse, from simple verbal reports, to eye tracking and button presses, to neural correlates such as EEG and fMRI. The influence of ‘top-down’ or motivational effects [13] can be readily examined in humans, who are easily instructed. In other animals, instruction is typically achieved by classical conditioning (associating neutral cues with punishment or reward). On the other hand, saliency-driven effects (e.g., brightness, loudness) are typically described as ‘bottom-up’ attention [13,14].

The ideal insect attention study would address all of the above criteria, using an experimental design capable of demonstrating unity, resource limitation, alternation, and neural correlates.

## Calcium imaging in behaving flies

To make the case for attention in the insect brain, we will begin with three recent *Drosophila* brain-imaging studies, even though these were not designed to query selective attention specifically. All three studies used two-photon calcium imaging of the brains of flies engaged in visually guided behaviors. Recent advances using genetically encoded calcium sensors in *Drosophila* provide a powerful new approach for uncovering attention-like processes in the insect brain, with the dynamic calcium signal being the resource of potential interest. In the first study by Aptekar et al [15], the authors use a tethered flight paradigm to identify lobula projection neurons that appear to be required for separating objects from their background - presumably a requisite step for paying attention to a specific object in the visual field. The anterior optic tract (AOT), where some of these neurons project, is especially implicated in figure/ground discrimination (Figure 1). Optic glomeruli in the wider lateral protocerebrum region containing the AOT could thus conceivably be where visual primitives (orientations, shapes, sizes) are encoded in parallel, as also suggested by an earlier electrophysiological study [16]. These studies therefore show evidence of parallel processing and visual filtering, which are prerequisites for attention [7], but no evidence of selection or alternation. The next likely relay station for visual stimuli appears to be the central complex (CX), so might selective processing occurring there?

**Figure 1.**
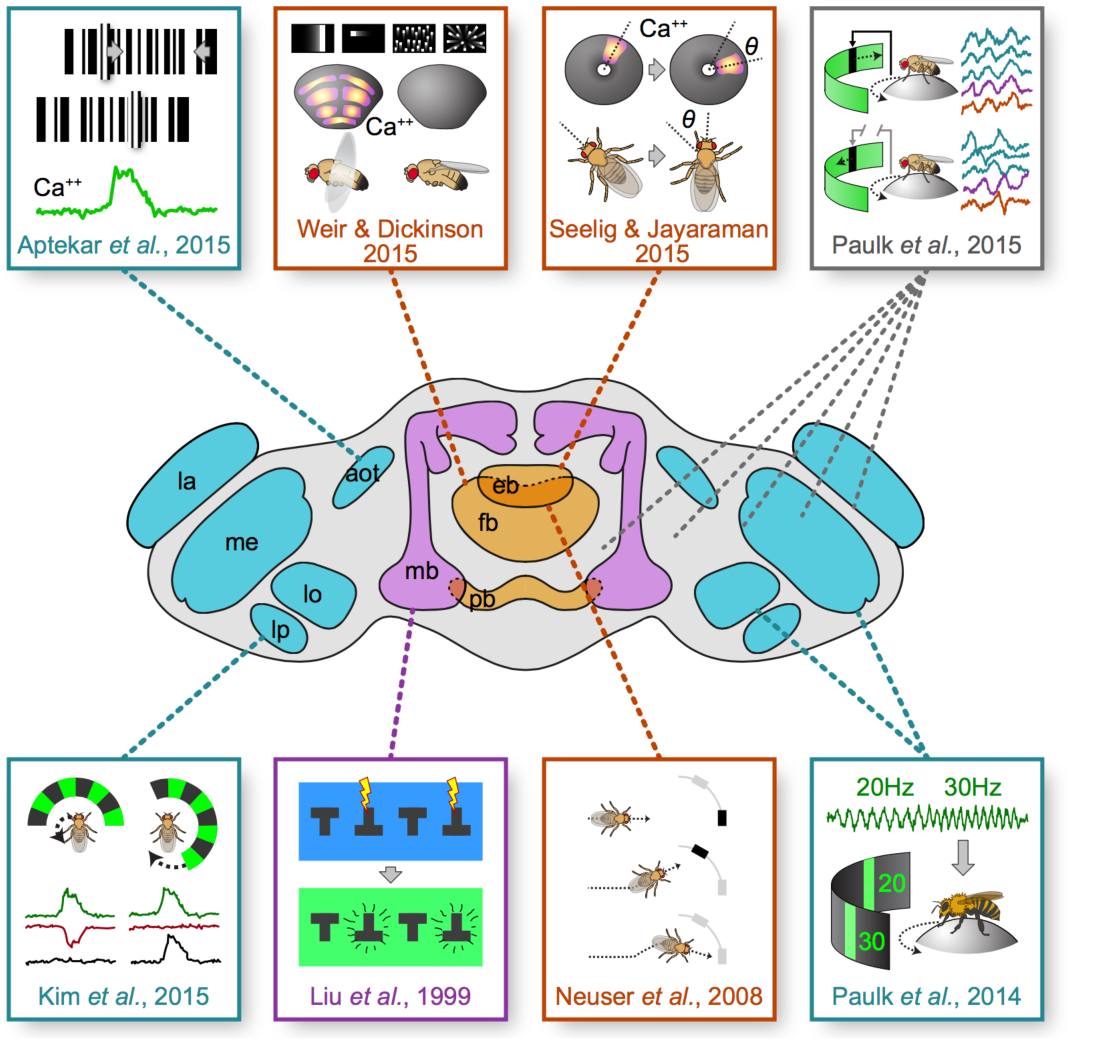
Attentional phenomena associated with specific brain regions. — Boxes illustrate the major findings of eight papers investigating attentional phenomena. Dashed lines connect findings to implicated neuropils. Aptekar *et al*. (2015) identified Ca++ transients correlated to figure-ground separation in the lobula (lo) and anterior optic tract (aot). Weir and Dickinson (2015) observed flight-gated tuning of modules in the fan-shaped body (fb) to visual stimuli. Seelig and Jayaraman (2015) discovered an “bump” of Ca++ activity in the ellipsoid body (eb) which tracks visual cues and locomotion in an egocentric frame. Paulk *et al*. (2015) observed synchronization in extracellular recordings across many brain regions, from the lamina (la) to the central brain, but only in animals engaged in closed-loop tasks. Kim *et al*. (2015) observed likely inhibitory efference copy (red trace) triggered by self-initiated motion in the lobula plate (lp). In an early study, Liu *et al*. (1999) showed that the mushroom bodies (mb) are required for context generalization during visual learning. Neuser *et al*. (2008) showed that the ellipsoid body is required for the maintenance of goal-oriented navigation in the presence of distractors. Lastly, Paulk *et al*. (2014) showed that extracellular potentials in the medulla (me) and lobula will synchronize alternatively with stimuli flickering at 20 or 30Hz presented to the ipsi-and contralateral visual fields. pb: protocerebral bridge.

In another recent *Drosophila* brain imaging study, Weir and Dickinson found that all structures of the CX respond to visual stimuli, although only the responses in the fan-shaped body (FB) depend on behavioral state (e.g., flight) [17]. Different layers within the FB respond to different objects and motion properties during flight, consistent with an earlier study assigning visual learning of different kinds of objects to different FB layers [18]. This suggests a level of filtering by FB layer, which therefore already provides a potential substrate for stimulus selection. Because of the columnar organization of the CX [19], activity in the FB is likely to be reflected by matched activity in other CX structures [19], such as the ellipsoid body (EB) and the protocerebral bridge (PB) - although it is interesting to note that visual responses in the EB were not gated by flight behavior. State-dependent modulation of circuit activity, and gain modulation [20,21] more generally, is likely an essential neural computation primitive supporting attentional processes. Another line of evidence pointing to the FB as a general arousal-setting center is its role in sleep/wake homeostasis [22]. This is telling, because sleep and attention might employ similar stimulus suppression mechanisms in the brain [23]. Certain layers of the FB (*e.g*, the sleep-inducing dorsal layer [24] might therefore be involved in the capacity to *suppress* information flow in the insect brain - but flow to where? The FB projects to the EB and the Lateral Accessory Lobes, making these areas plausible destinations of filtered information [19].

A third recent imaging study in *Drosophila* by Seelig and Jayaraman looked at EB activity in walking flies [25]. The EB contains concentric ring neurons, wedge neurons, and tile neurons [26], assuming a donut shape in flies or a croissant in some other arthropods [27]. The authors imaged calcium activity in the wedge neurons of the EB, as tethered flies oriented towards vertical bars in a closed-loop virtual reality environment. This peek into the behaving fly’s brain led to arguably the most convincing evidence so far for a neural correlate of visual attention in the insect brain: a discrete ‘bump’ of activity mapped onto the EB wedges, which seemed to correspond to the angular position of the fixated object. This discovery is reminiscent of the discovery of ‘place cells’ in rodents [28], not as much for the specific processes involved as for the sheer potential this activity feature now provides for understanding information processing in the brain.

Several aspects of the EB activity ‘bump’ suggest it reflects an attention-like phenomenon. First, the bump is egocentric: while its position in the EB maps accurately onto the azimuthal position of the fixated object, this mapping can be completely different across individuals. Second, when presented with multiple objects to fixate on, the bump remains a single unitary bump. Third, the absolute activity level of the bump appears to fade in and out, as if other sensory processes are competing for this limited resource. Fourth, when flies are presented with multiple objects, the bump can jump to an alternate section of the EB, consistent with serial alternation. Finally, the activity bump persists in animals that are not walking, and can even persist for several seconds in complete darkness. This last observation supports an earlier *Drosophila* study that showed the EB is required for maintaining fixation on temporarily invisible objects [29].

It would appear that Seelig and Jayaraman may have identified a neural correlate for goal-directed attention in the insect brain. It remains to be shown whether this egocentric map generalizes to other behaviors (e.g., flight) or sensory modalities (e.g., olfaction), or if it is modulated by some behavioral states (e.g., sleep). Importantly, the three imaging studies highlighted above uncover different features of visual filtering in the insect brain, from figure/ground discrimination in optic glomeruli [15] to state-dependent sorting of visual primitives in the FB [17] to selecting (and ignoring) competing objects in the EB [25]. However, filtering alone is insufficient for selective attention; at some level a binary decision is made in order to promote unambiguous behavioral outputs. How discrete activity patterns the EB might lead to selective motor output will be a likely focus of future *Drosophila* attention work.

## Earlier behavioral evidence from visual learning studies

Before the current spate of *Drosophila* brain imaging studies, the best evidence for attention-like processes in the insect brain pointed to the mushroom bodies (MB) (Figure 1), in visual learning studies in flying *Drosophila* [30-33]. While these studies were also not specifically designed to measure attention processes [14], the common conclusion from most of this work is that the MB are required for separating the visual context (e.g., background color or texture) from the object that must be remembered. This filtering mechanism resembles figure/ground discrimination specifically and selective attention generally, in that the context must be actively ignored. Without the MB, flies fail to ignore the non-predictive context in visual learning assays [30]. Additionally, the MB, and dopaminergic input to the MB, seems to be required for regulating visual salience and decision-making [32]. Typically considered an olfactory learning and memory neuropil in dipteran species such as *Drosophila*, the MB also receives inputs from higher-order visual centers through the accessory calyx in bees [34] and may process visual cues similarly to olfactory cues in flies [35]. The MB also seems to be required to maintain behavioral flexibility and to delay motor learning [36], which is probably important for maintaining a window for selective attention to operate before habit formation sets in.

Just as the MB appears to mediate both learning and attentional processes, the EB, which contains the attention-like bump, is required for visual and spatial learning behaviors [37,38]. An appealing hypothesis is that the same molecular and synaptic processes that mediate learning can facilitate attention-like processes [39,40]. In vertebrate systems, selective attention probably involves coordinated activity across the brain rather than the output of a specific set of neurons [41], so it is perhaps not surprising that for attention-like processes in insects, a variety of learning and memory circuits might be implicated (Figure 1). Perhaps similarly, sleep regulation has also been assigned to a variety of circuits, often the same CX and MB structures uncovered from visual learning studies [24,42]. If selective attention involves the recruitment of disparate neurons into transient ensembles, as it appears to do in the mammalian brain [41], then focus on any one structure might not be really informative.

Nevertheless, the most convincing evidence to date has been found in the central complex, and specifically the EB [25]. One potential explanation for why the central complex might contain conspicuous neural correlates of selective attention relates to its role in controlling locomotion [43]. Selective attention most likely emerged to help motile creatures anticipate (or avoid) events as they moved through a cluttered environment [44]. There may therefore be a deep link between selective attention and locomotion control mechanisms: coordinated activity among central complex modules [19] (e.g., the aforementioned wedges of the EB) may guide locomotion, and concomitantly provide a ready architecture for more sophisticated attention-like processes.

## Explicit tests of behavioral attention

Several bee studies have been specifically designed to identify and measure visual attention behaviors [6,9,11,45,46], many of these inspired by human attention experiments. The critical problem for any behavioral attention study, solved by bee researchers long ago, is how to get an insect to do the equivalent of ‘pushing a button’. Honeybees (*Apis mellifera*) will readily visit visual objects that have been associated with a reward, such as sugar water, so it is relatively easy to design experiments to measure distractor effects or reaction times - key measures for attention experiments (Figure 2). For example, using such free-flight assays, one study found that honeybees engage in a narrow serial search for visual targets, whereas bumblebees (*Bombus terrestris*) seem to perform a broader parallel search [9]. This last result in bumblebees demonstrates a capacity for feature-based attention in insects, whereby disparate objects grouped by a commonly attended feature (e.g., a color) are easier to detect [47].

**Figure 2.**
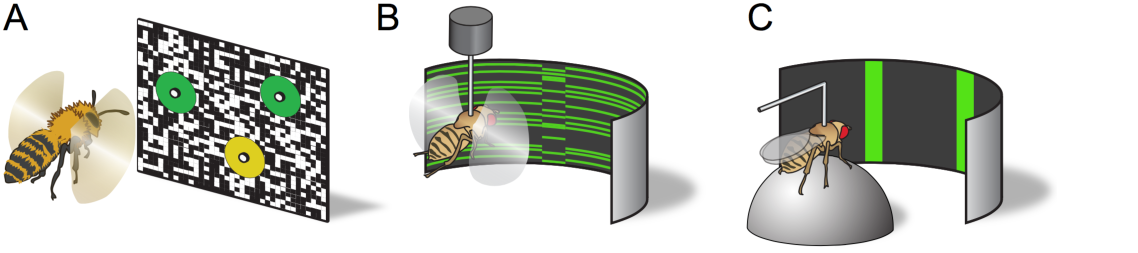
Versatile experimental assays for investigating attention. — A) Bees trained to feed at a particular site (the yellow circle) choose among competing sites (green circles) while being challenged by visuals distractors (in this case, the high contrast random dot pattern). B) Measured torques or wing-beat amplitudes from tethered flying flies can close a feedback loop on visual stimuli, in this case a figure-ground stimulus. C) As in B, but with a tethered fly walking on an air-supported ball. The visual stimulus here represents competing objects between which the fly’s fixation can alternate.

Compared to *Drosophila*, honeybees lack genetically encoded reagents permitting pan-neuronal or targeted physiological imaging, but are well suited to free-flight visual choice experiments, because such paradigms exploit their innate visually guided foraging strategies. Designing free-flight assays to probe attentional processes in flies is considerably harder (but see [48]). Instead, the only way that selective attention has been directly investigated in flying dipterans so far is using tethered paradigms (Figure 2). This approach does offer advantages: the visual context can be carefully controlled and behavioral readouts can be tracked on a millisecond timescale. Early studies by Heisenberg and Wolf showed that *Drosophila* flies alternate between competing moving objects, such that they can suppress responses to one while they fixate in closed loop on the other [49]. In more recent work, Sareen *et al*. used LED arenas to show that competing optomotor responses in open loop can be biased to the left or the right by a preceding visual cue (a flashing light), suggesting that flies have a dynamic ‘focus of attention’ [8]. Importantly, cueing effects could be separated in time (by up to 4 seconds) from the visual attention test, suggesting a characteristic timescale (Box 1) for the focus of attention in wild-type flies. Recent follow-up work confirms the surprising finding that wild-type flies indeed have a measurable ‘attention span’ of about four seconds in this paradigm [10].

## Evidence from electrophysiology

In the visual experiments described above, the object of a fly’s attention was inferred by flight bias toward one side, in the face of competing objects on either side [8,10]. While behavioral readouts are crucial for any attention experiment in animals other than verbal humans, electrophysiology provides important supporting evidence for stimulus selection and suppression in the brain, especially for the faster temporal readouts characteristic of selective attention, such as neural oscillations [50]. Tang and Juusola [51] performed a similar experiment as Sareen et al (minus the cuing effects) in tethered flies implanted with a recording electrode in each optic lobe. The authors reported increased neuronal spiking and local field potential (LFP) activity on the side facing the fixated cue (and suppressed activity in the other optic lobe). Hinting at seriality, brain activity changes alternated between the competing stimuli (moving gratings presented to either eye), and these changes even seemed to precede the behavioral choices. LFPs were in the 20-50Hz range, consistent with previous studies that found LFP signatures for visual salience and novelty in the *Drosophila* brain [39,40,52-54]. Together, these studies suggest an endogenous oscillation in the insect brain associated with selective attention, perhaps similar to alpha (8-12 Hz) or gamma (30-90 Hz) oscillations that have been linked to attention processes in mammals [50]. It will be interesting to determine in future studies where in the insect brain these oscillations originate (but see [55] for evidence in crayfish, which suggests the oscillations are widespread), and how they might be employed for information processing.

A complementary approach to tracking neural correlates of attention in the brain is to impose oscillations exogenously using periodic stimuli. Thus, contrasting visual flickers (e.g., one object flickering at 10Hz and another at 15Hz) will evoke overlapping oscillatory activity throughout the insect brain, which can be disambiguated and contrasted in the frequency domain. Such experiments, long a staple of human attention research [56], have only recently been applied to flying and walking insects in visual closed-loop visual attention paradigms [57,58] (Figure 2). One conclusion from these studies, consistent with human attention studies, is that fixation on a flickering object increases the amplitude of that oscillation in the brain, at the expense of competing flickers which are suppressed, even in early layers of visual processing [57]. This suggests that neural selection and suppression mechanisms characteristic of selective attention are not necessarily confined to central brain structures such as the EB, FB, or MB. They might already operate in the periphery, consistent with experiments showing that behavioral states modulate neural activity in the optic lobes [20,21].

Support for attention-like processes in the optic lobes has been found in other species as well. Weiderman and O’Carroll recorded spiking activity from centrifugal neurons (small target motion detectors) in the optic lobes of dragonflies and found unitary responses when the insects were presented with multiple competing target objects [59]. While these responses were still linked to specific retinal receptive fields (i.e., they were not egocentric, like the ‘bump’ in the EB), which response ‘won’ or ‘lost’ seemed to be under the control of other circuits, perhaps in the central brain. While these dragonfly experiments did not include any behavioral readout, one prediction is that these neural events might represent a behavioral choice (e.g., what target might the dragonfly have pursued?). Recent behavioral and theoretical work on dragonfly prey capture suggests that, like mammals, insects rely on internal models to guide their actions [60,61], so it is possible that selective suppression of visual responses in the insect optic lobes reflects output from these centralized models.

A conceptually similar result was recently reported in a *Drosophila* study, which combined electrophysiology and behavior (tethered flight torques in a visual arena) [62]. Kim *et al* found that activity in motion-sensitive lobula neurons (evoked by moving gratings) was suppressed by ‘voluntary’ flights to the opposite side (Figure 1). This suggests that motor decisions can transiently attenuate neural responses to conflicting sensory stimuli, for example to preventing a conflict between optomotor reflexes [63] and self-initiated behavior. Interestingly, conflicting flight decisions do not attenuate lateralized calcium signals in the FB [17], suggesting that a different level of filtering is occurring in the central brain.

In conclusion, accumulating evidence from a number of studies supports the view that insects have selective attention, at least for vision (and see [64,65] for evidence on auditory attention in insects). What seems less certain is which brain regions or neurons might be involved regulating attention-like processes. So far, we have discussed a potential role for almost every neuropil in the insect brain (Figure 1), except the outermost layers of the optic lobes. One way to explain these varied observations is that successive visual processing layers (such as figure/ground discrimination, followed by state-dependence filtering) feed into attention centers, which might be more localized in the central brain. An alternative view might be that attention is not confined to any one structure, but is instead a brain-wide phenomenon whereby far-flung neurons are recruited into functional ensembles, yielding unitary outcomes [41]. This hypothesis presents a serious experimental and conceptual challenge, especially for researchers using model systems such as *Drosophila* who may be accustomed to a toolkit of reductionist reagents, such as sparsely expressed transgenic drivers. We suggest that to understand attention-like processes in the insect brain (or any brain), one must ideally manipulate and/or record from disparate neurons or structures simultaneously. Future research using whole brain 4D volumetric imaging [66,67] or multichannel electrophysiology [54,68] in behaving animals might be a viable option.

## Conflict of Interest

The authors have no financial conflicts of interest.

## Acknowledgements

BldB was supported by a Sloan Research Fellowship. BvS was supported by an Australian Research Council Discovery Project DP140103184.

